# Understanding early reproductive failure in turtles and tortoises

**DOI:** 10.1101/2023.06.07.544015

**Authors:** Alessia M Lavigne, Robert Bullock, Nirmal Jivan Shah, Chris Tagg, Anna Zora, Nicola Hemmings

**Author notes:** Article impact statement: Informing conservation with a novel and practical solution for identifying causes of early reproductive failure in turtles and tortoises.

## Abstract

Turtles and tortoises (Order Testudines) are facing an extinction crisis, and ecosystems are at risk of collapsing with the loss of key roles they play. Hatching failure is a crucial barrier to population growth and persistence, but its causes are poorly understood, and it is unknown whether fertilisation rates are declining as many populations become smaller and more female-biased. Here, we first show that fertilisation rates are considered in only a very small proportion of studies of turtle and tortoise hatching success, and those studies that do attempt to measure fertilisation rates use unreliable methods. We also show that studies of hatching success are biased towards marine turtles, as opposed to freshwater and terrestrial species, and wild rather than captive populations. We address the lack of reliable methods for assessing fertilisation rates in turtles and tortoises by developing and testing a microscopy-based method for detecting perivitelline membrane (PVM) bound sperm and embryonic nuclei in the germinal disc of unhatched eggs. We demonstrate that this method can provide unequivocal evidence of egg fertilisation in three different turtle and tortoise species from both captive and wild populations, even after eggs have been left in the nest for the full incubation period. This approach therefore represents a valuable new tool for monitoring egg fertility and embryo survival rates in turtles and tortoises, with the potential to provide important insights into the underlying drivers of reproductive failure in threatened populations.

## Introduction

Turtles and tortoises (Order Testudines) are facing severe population declines: over 50% of species are threatened (includes all critically endangered, endangered, and vulnerable species), of which 20% are critically endangered (Lovich *et al*., 2018; Rhodin *et al*., 2018; Stanford *et al*., 2020). Since 1800, at least seven species have gone extinct, three of which have been lost in the past few decades (Stanford *et al*., 2020). Many threats faced by turtles and tortoises impact their reproductive success and lead to reduced hatching success (Lovich *et al*., 2018). Low hatching rates have been reported for several threatened species, including the Leatherback turtle (*Dermochelys coriacea*) (50.4%; Rafferty *et al*., 2011), Olive Ridley turtle (*Lepidochelys olivacea*) (8%; Honarvar *et al*., 2008), and green turtle (*Chelonia mydas*) (20-60%; Booth *et al*., 2022), and are predicted to decline further under climate change in several species (Fuentes *et al*., 2010; Pike, 2014).

Several factors may impact hatching rates in wild turtle and tortoise species. Biotic drivers include low egg fertilisation rates, potentially linked to male/sperm availability (Miller, 1985); genetically determined developmental abnormalities in embryos (Ingle *et al*., 2021); microbial/fungal infection of eggs (Peters *et al*., 1994; Gleason *et al*., 2020; McMaken, 2022; Carranco *et al*. 2022); maternal condition (Rafferty *et al*., 2011; Duchak and Burke, 2022); and the density of females at a nesting site (Booth *et al*., 2022). Habitat fragmentation may also indirectly lead to reduced hatching success and reproductive issues, via inbreeding depression and low genetic diversity (Ennen *et al*., 2010). Hatching success may also be influenced by abiotic factors such as rainfall, flooding, substrate composition and water potential, temperature, salinity, and pollution (Ragotzkie, 1959; Mortimer, 1990; Wood and Bjorndal, 2000; Bilinski *et al*., 2001; Stanford *et al.,* 2020; Limpus *et al*., 2021), as well as elevation, slope, and erosion of nesting beaches (Kraemer and Bell, 1980; Maneja *et al*., 2021).

Hatching failure is also a problem for captive turtle and tortoise populations. Captive conditions can negatively affect reproductive condition, potentially leading to male infertility or behavioural/copulation issues, low fertilisation rates and decreased hatching success (Currylow *et al*. 2017), as well as low levels of reproductive hormones (Bin *et al*., 2010). In addition, translocation may stifle breeding success for several years (Currylow *et al*. 2017). Hatching rates in captive marine turtles have been shown to be consistently lower than in the wild (Owens and Blanvillain, 2013), perhaps due to fertility issues, or captivity-related impacts on embryo survival (e.g., lack of essential fatty acids in maternal diet; Craven *et al*., 2008; Owens and Blanvillain, 2013). However, in Gopher tortoises (*Gopherus polyphemus*), hatching success was found to be higher in captivity (58.8%) than in natural nests (16.7%), suggesting that low hatching success in the wild is attributable to both intrinsic (egg quality) and extrinsic (nest environment) factors (Noel *et al*., 2012). Nevertheless, many captive breeding programmes are plagued by elevated hatching failure. For instance, the sole remaining female Yangtze giant softshell turtle (*Rafetus swinhoei*) produced undeveloped eggs for over eight years before her death in 2019, despite international conservation efforts, including artificial insemination (Lovich *et al*., 2018; Liu *et al.,* 2019).

Identifying the underlying reproductive barriers to hatching success is crucial for improving success rates of both wild and captive conservation efforts (Bell *et al*., 2003; Phillott and Godfrey, 2020; Dovč *et al*., 2021). Hatching failure can result from either fertilisation failure or embryo death, and these two issues may have different causes (Hemmings *et al.,* 2012). Unfertilised eggs are indicative of parental fertility, copulation, or behaviour problems, whereas embryo mortality is more likely due to genetic or developmental problems and/or environmental factors directly affecting the egg or embryo (Hemmings *et al.,* 2012).

Monitoring fertilisation rates also serves a broader goal of understanding the impacts of environmental change on turtles and tortoises (Order Testudines). Eggs are typically deposited by the mother in burrowed nests and left to incubate at ambient temperatures, and the effect of global warming on incubation temperature is therefore one of the most significant conservation concerns for many species (Stanford *et al*., 2020). With the current rate of global warming, incubation temperatures are likely to exceed the range of 25°C – 34°C required for viable embryo development in populations close to the equator (Ackerman, 1997; Hays et al., 2017; Hawkes *et al*., 2007; Hays *et al*., 2022). Even if embryos survive high incubation temperatures, many turtle and tortoise species have temperature-dependent sex determination, where embryos usually develop into males at cooler incubation temperatures and females at warmer incubation temperatures (Ewert *et al*., 2004; Valverde et al., 2010). A temperature-induced female bias at the population level could lead to a reduced egg fertilisation rate, loss of genetic variation, and decreased effective population size (Montero *et al.,* 2018; Hays *et al*., 2022). These issues have already been identified in certain populations and require urgent monitoring and intervention if they are to be resolved (Hawkes *et al*., 2007; Fuentes *et al*., 2010; Booth *et al*., 2020; Chatting et al., 2021; Hays *et al*., 2022). Monitoring rates of fertilisation failure and embryo death may provide an early indicator of population sex bias (e.g., more unfertilised eggs due to insufficient males/sperm) or high levels of temperature-related embryo mortality, allowing more rapid conservation intervention.

Conservation interventions may themselves also influence hatching success (Marshall *et al*. 2023). For example, nest relocations are commonly used to reduce threats such as tidal inundation (e.g., McElroy et al., 2015), but data suggests that manipulation of eggs may contribute to egg failure through increased embryonic death (Wyneken et al., 1988). The impact of nest relocation on hatching success is unclear: some studies report lower hatching success compared to undisturbed nests (e.g., Garrett et al., 2010; Candan, 2018), while others show improvements provided sites are carefully chosen (e.g., Wyneken et al., 1988). Measuring the proportion of embryo death in relocated nests is therefore a useful tool for conservation managers, allowing them to assess how successful or disruptive relocation attempts are, and to measure the suitability of different nest relocation sites.

Despite the importance of distinguishing between fertilisation failure and embryo survival as causes of hatching failure, few studies of turtle and tortoise eggs appear to do so. Undeveloped eggs with no visible embryo may be unfertilised, or they may contain an early-stage embryo that died before it was visible to the naked eye. Embryonic development begins within the mother’s oviduct prior to oviposition, so by the time a fertilised egg is laid, the developing embryo is already several days old. This means that embryo death can even occur prior to oviposition (Abella *et al*., 2017). In published studies to date, undeveloped eggs are typically classified as unfertilised without further examination (e.g., Langer *et al*., 2020; Gane *et al*., 2020a), and where attempts are made to determine fertilisation success, potentially inaccurate macroscopic methods are typically used (Gárriz *et al*., 2020; Phillott and Godfrey, 2020). Recently, a small number of reptilian captive breeding programmes have trialled microscopic techniques, originally developed for birds (Birkhead *et al*., 2008; Hemmings *et al*., 2012), to help investigate infertility (Croyle *et al*., 2015; Croyle *et al*., 2016; Augustine, 2017), and the application of these methods has also been recommended for assessing fertility of wild sea turtle eggs (Phillott and Godfrey, 2020; Phillott*, et al*., 2021). The techniques allow detection of sperm on the perivitelline membrane (PVM) surrounding the yolk and embryonic nuclei in the germinal disc, thereby providing unequivocal evidence of fertilisation. In birds, these techniques have revealed that approximately 52% of unhatched eggs are misclassified as ‘unfertilised’ using traditional macroscopic techniques (Hemmings and Evans, 2020). However, reptilian studies have so far only been successful in identifying PVM-bound sperm in freshly laid, unincubated eggs. The ability to identify embryonic nuclei in unhatched eggs has not been demonstrated, and the methods have not been tested on eggs that have been incubated. To be useful for in situ conservation, these techniques must be applicable to unhatched eggs recovered from nests at the end of the incubation period (Phillott and Godfrey, 2020), which have often experienced decomposition, desiccation, and insect infestations (Abella *et al*., 2017).

The aims of this study are two-fold. In Part 1, we review the existing literature on turtle and tortoise hatching success to determine the extent to which studies differentiate between fertilisation failure and embryo mortality as causes of hatching failure and how this varies across taxa. We also critically assess the methods currently used for examining failed eggs. In Part 2, we develop and test a method for detecting perivitelline-bound sperm and embryonic nuclei in the germinal disc of unhatched eggs from three captive species, the Red-footed tortoise (*Chelonoidis carbonarius*), Galapagos giant tortoise (*Chelonoidis nigra*), and Spiny turtle (*Heosemys spinosa*), and one wild species, the Hawksbill turtle (*Eretmochelys imbricata*), collected at the end of the incubation period.

## Part 1: Systematic review of turtle hatching success literature

### Part 1: Methods

We searched English abstracts from papers on Web of Science Core Collection and Scopus on 24 November 2021 using the following terms: (“hatching failure” OR “hatching success” OR “hatchability” OR “hatching rate”) AND (“turtle” OR “tortoise” OR “testudines” OR “chelonia”). Our search may not have retrieved every article published on this topic, but since the chosen databases are considered world-leading (Zhu and Liu, 2020), we assumed the retrieved articles represented most of the relevant published literature. Combined, the databases returned 374 records, but we were unable to include 44 papers (6.7%) due to lack of institutional access (23 papers) and/or because they were not published in English (21 papers; Figure 1). We read the final set of papers in detail to determine (a) the methods used to determine egg fertility status, (b) the species studied, and (c) whether the population was wild or captive.

**Figure 1.**
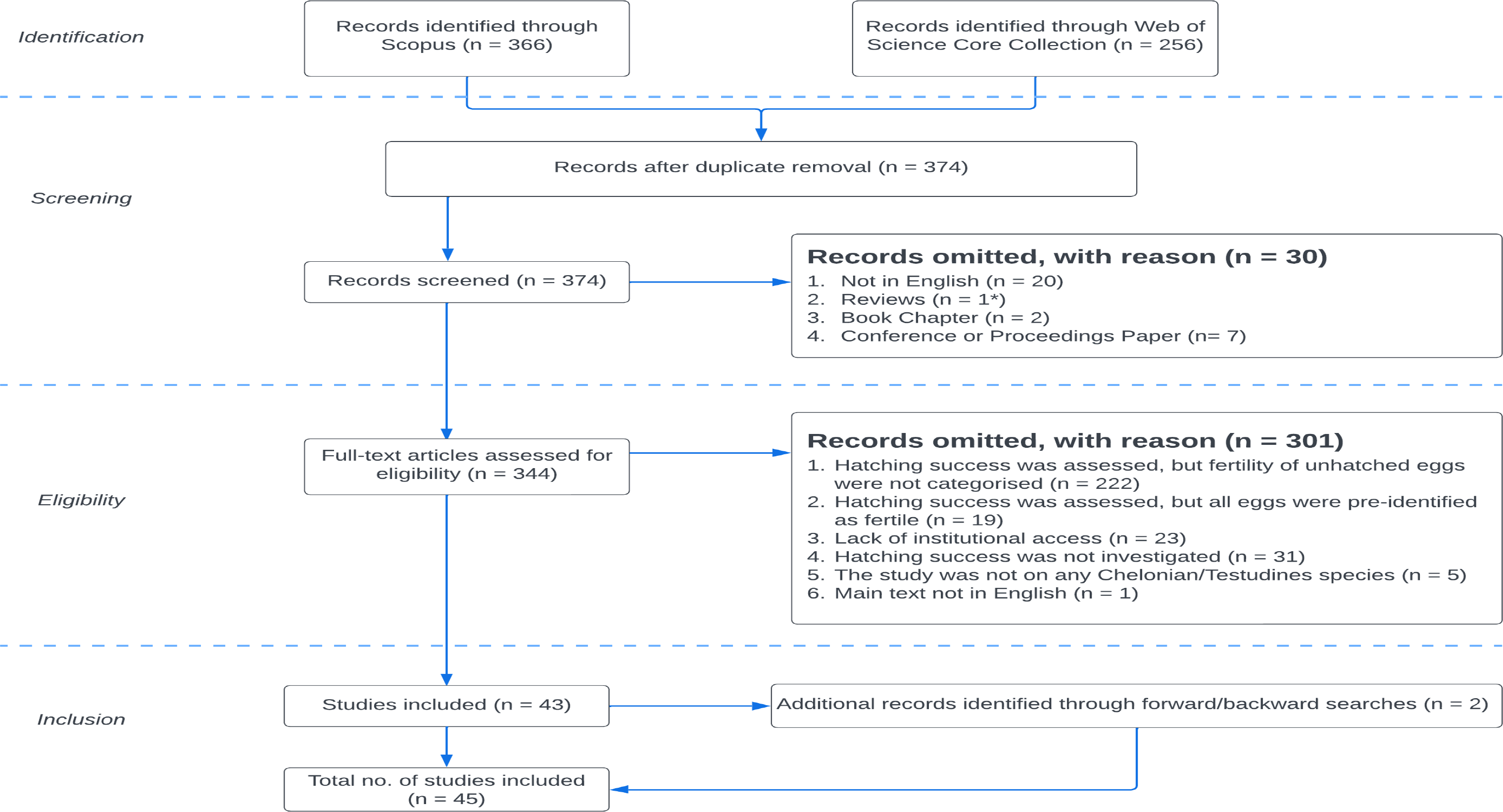
*Depiction of the literature search and screening process to filter English studies that have claimed to investigate both Chelonian/Testudines hatching success and fertility of unhatched eggs, following Preferred Reporting Items for Systematic Reviews and Meta-Analyses (PRISMA) (**Moher* et al., *2009;* *O’Dea* et al., *2021**). Forward/backward searching involved identifying and examining relevant papers cited in an eligible article or relevant papers that cite back to a specific eligible article.* *\*The original number of records classified as “Reviews” was n = 4, but upon inspection, three were incorrectly categorised and were primary research articles, and therefore included for the ‘Eligibility’ stage.*

### Part 1: Results

#### Number and proportion of studies assessing egg fertility

We identified 286 studies that investigated turtle and tortoise hatching success (Figure 2A: includes records omitted during eligibility checks under [1]: n = 222 and [2]: n = 19, as well as total number of studies included, n = 45, from Figure 1). Of these, 76.62% (n = 222) did not assess the fertility status of unhatched eggs, and 6.64% (n = 19) specified that they worked with eggs with visible signs of development only (Figure 2A). Only 15.73% (n = 45) attempted to classify the fertility of unhatched eggs (see Appendix A for additional information on methods, species, and population context for these studies). Most studies (∼84%) investigating hatching success have therefore overlooked the potential role that variation in egg fertility may play.

**Figure 2.**
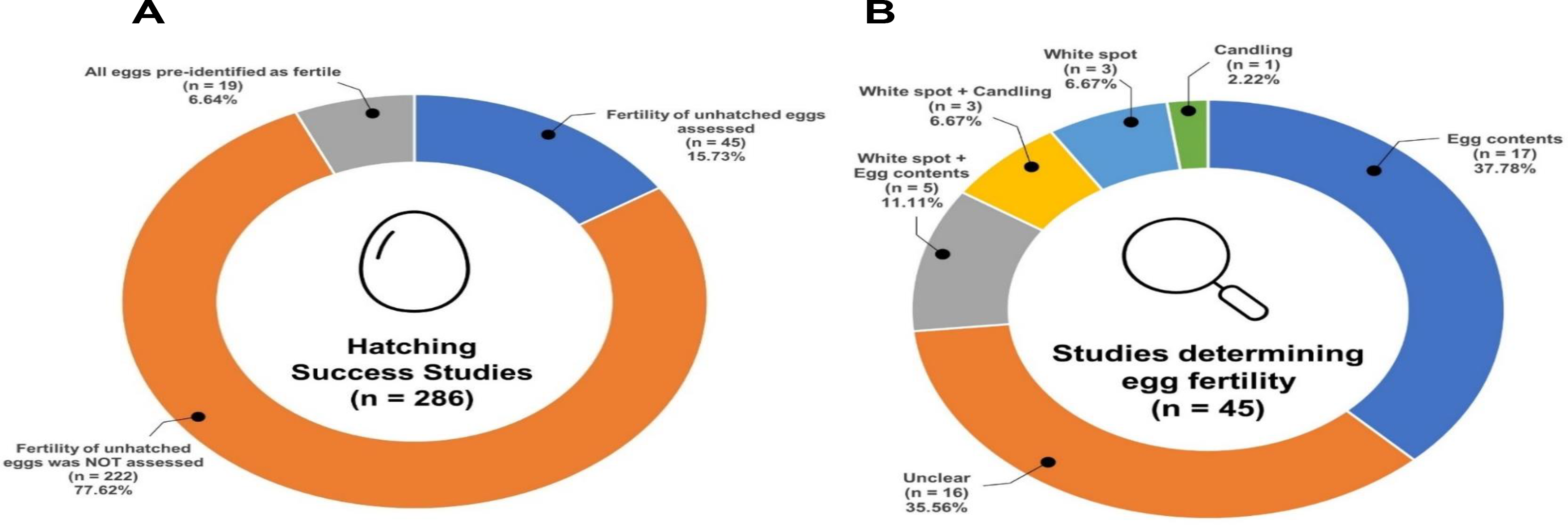
*Number and proportion of papers on turtle and tortoise (Order Testudines) hatching success (A) that assessed egg fertility. (B) categorised by their chosen methods of determining egg fertility.*

#### Methods used to determine fertility status of eggs

We found that studies used a range of different methods to assess fertilisation status of turtle and tortoise eggs (Figure 2B). The majority (37.8%) used macroscopic visual assessment of egg contents to determine the fertility of unhatched eggs, which typically involves examination of egg contents for visible blood or embryonic tissue (Wyneken *et al*., 1988) and overlooks early-stage embryos (Hemmings & Evans, 2020). In some studies, all undeveloped eggs were explicitly categorised as “infertile” (Ulrich and Parkes, 1978) despite not being examined further. This should be avoided to prevent misunderstanding of the underlying reproductive problem.

A smaller proportion of studies based their fertility assessments on the presence of a white spot on the eggshell and/or visibility of embryo development on candling (Figure 2B). Unlike bird eggs, reptile eggs do not possess a chalaza (strands of albuminous matter that position the yolk and developing bird embryo in the centre of the egg), so the yolk, which has a lower density than the surrounding albumen, floats to the top of the egg and the embryo develops next to the shell membrane. This process occurs soon after oviposition and proceeds with the shell membrane adhering to the shell, creating a white spot on its surface, often referred to as “chalking” (Blanck and Sawyer, 1981; Phillott and Parmenter, 2007; Matsubara *et al*., 2016; Phillott and Godfrey, 2020). Eggs without a white spot are therefore often interpreted as being unfertilised. However, the visibility of the white spot varies (Phillott and Godfrey, 2020) and it can disappear within ∼44 hours of embryonic death (Phillott and Parmenter, 2007). Consequently, eggs that experience early embryo death several days before collection can be misclassified. Eggs without a white spot therefore require further inspection before being classified as unfertilised (Phillott and Godfrey, 2020). Despite this, we found that white spot presence was used to determine egg fertility in 24.4% (n = 11) of studies in our search and was the sole indicator in 6.7% (n = 3).

Four studies (8.89%; Figure 2B) used candling to assess fertility status, which involves shining a bright light through the eggshell, allowing the observer to identify embryonic development and desiccation (Dovč *et al*., 2021) without opening the egg. Candling is a quick, non-invasive, and useful method, but as with white spot assessment, it is not completely reliable for determining whether an egg is fertilised. Abella *et al*. (2017) was able to identify embryonic development as early as 24 hours post-oviposition via candling, but it remained unclear whether eggs that did not show obvious development were unfertilised. Via macroscopic assessment of egg contents, Dovč *et al*. (2021) showed that some eggs determined as “unfertilised” by candling were indeed fertilised.

Finally, more than a third of studies (35.56%) did not explain the methods they used in enough detail to be categorised (Figure 2B; “unclear”, also see Appendix A).

#### Taxonomic bias

Our final set of 45 studies (Figure 2B) focused on 23 species, representing just ∼7% of all extant Testudines (Figure 3). Marine turtles were the best studied, with 5 of 7 marine turtles represented and at a much higher frequency than any other group (30 investigations of marine turtles in total, exceeding the number of freshwater turtle and tortoise studies (n = 23) combined; Figure 3). The Loggerhead Turtle (*Caretta caretta*) was the most investigated species (13/45 studies (29%)). Freshwater turtles have the greatest number of species studied (total of 17 species), but at a relatively low intensity (≤2 studies per species). Tortoises are under-researched – represented by a single study only, on the Ploughshare Tortoise (*Geochelone yniphora*; Appendix A; Bourou *et al*., 2001).

**Figure 3.**
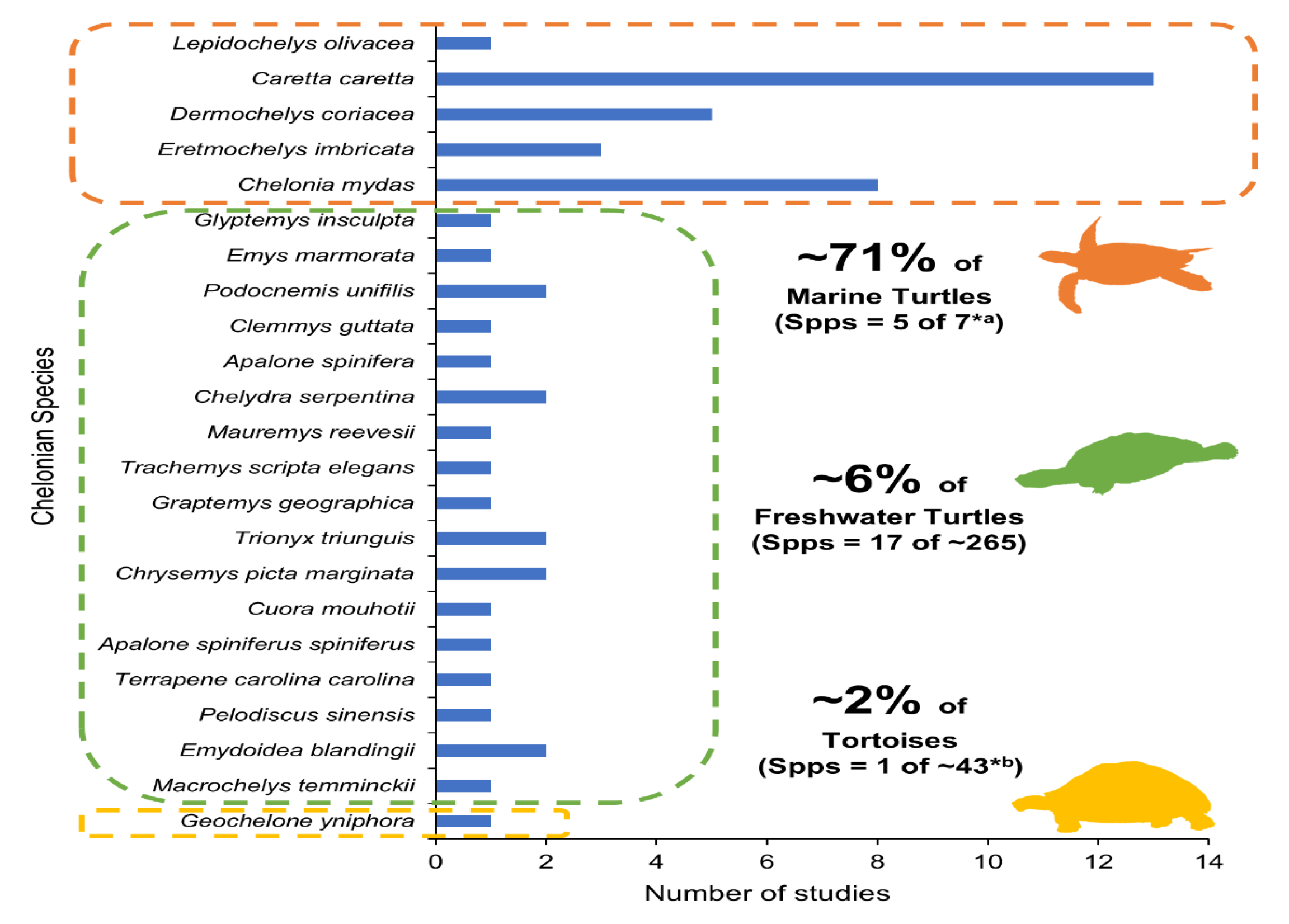
*The taxonomic distribution of papers reporting on turtle and tortoise (Order Testudines) egg fertility. Data are from 45 papers in total, some of which report on more than one species. 23 species have been studied in total; thus, approximately 292 (∼93%) of turtle and tortoise species have yet to be investigated. Determining approximation of total number of extant turtle and tortoise species: **\*^a^**Marine turtles: n = 7 (Rhodin* et al., *2018); **\*^b^** Tortoises (i.e., terrestrial turtles): n = ∼43 (Vlachos and Rabi, 2018); Freshwater turtles: Total number of extant species = ∼315 (Pough, 2013) – (*a + *b) = ∼265*

#### Population context bias: wild versus captive

We found a strong bias towards the study of eggs collected from wild clutches (84.44%), and few studies examining eggs from captive populations (13.3%), or both captive and wild populations (2.22%).

## Part 2: Detecting perivitelline-bound sperm and embryonic nuclei in the germinal disc of unhatched turtle eggs

### Part 2: Methods

#### Egg Collection and storage

We obtained 45 eggs from captive populations of Red-footed tortoise *Chelonoidis carbonarius,* Galapagos giant tortoise *Chelonoidis nigra*, and Spiny turtle*, Heosemys spinosa* via British and Irish Association of Zoos and Aquariums (BIAZA) UK Zoo members (Crocodiles of the World, Oxfordshire, and ZSL Whipsnade Zoo). Captive unhatched eggs were removed from incubators as part of standard zoo management procedures and transported to The University of Sheffield, UK, where they were refrigerated or frozen prior to dissection. All eggs were first examined for ‘traditional’ indicators of fertilisation commonly used in the existing literature (see Part 1). Eggs were considered “fertilised” according to traditional indicators if eggshell chalking/white spots were observed or visible embryo development was seen in the egg contents (Wyneken *et al*., 1988; Dovč *et al*., 2021). Eggs that displayed blood spots but no other signs of development were examined further to ascertain egg fertility (Table 1). Of the 45 captive eggs received, 27 showed no sign of development or had a blood spot only and were therefore examined microscopically (see below).

**Table 1.** *84 undeveloped turtle and tortoise eggs from four species were examined microscopically. 80 eggs showed no signs of chalking or embryo development. 4 had blood spots and these were also examined to determine whether they were fertilised as blood spots may be of ovarian origin (all 4 eggs were successfully examined). All wild eggs are from Hawksbill turtle populations in the Seychelles; all other eggs are from captive populations in UK zoos. All eggs were dissected for examination of the perivitelline membrane (PVM) and/or germinal disc (GD), but not all had retrievable or usable PVM/GD, resulting in a total of 67 eggs successfully examined.*

| Species | Common name | Conservation status | Population context | Storage prior to dissection |  |  | N° of undeveloped eggs | N° of eggs with blood spot | N° of eggs examined |
| --- | --- | --- | --- | --- | --- | --- | --- | --- | --- |
|  |  |  |  | Refrigerated | Frozen | Formalin-fixed |  |  |  |
| *Chelonoidis carbonarius* | Red-footed tortoise | NE | Captive | 1 | 2 | 0 | 2 | 1 | 3 |
| *Chelonoidis nigra* | Galapagos giant tortoise | NE | Captive | 2 | 21 | 0 | 20 | 3 | 21 |
| *Heosemys spinosa* | Spiny turtle | EN | Captive | 1 | 0 | 0 | 1 | 0 | 1 |
| *Eretmochelys imbricata* | Hawksbill turtle | CR | Wild | 0 | 0 | 58 | 58 | 0 | 42 |
| **Total** |  |  |  | **4** | **22** | **58** | **80** | **4** | **67** |

We also obtained 58 Hawksbill turtle eggs from wild populations from the Republic of Seychelles (hereafter referred to as Seychelles) during the 2021/2022 nesting season (Table 1), supported by partnerships with conservation organisations on three islands: Nature Seychelles (Cousin Island: 4.3315°S, 55.6620°E), Save Our Seas Foundation: D’Arros Research Centre (D’Arros Island: 5.4180°S, 53.2962°E) and Frégate Island Foundation (Frégate Island: 4.5837° S, 55.9386° E). Seychelles has one of the five largest Hawksbill turtle populations (Hitchins *et al.,* 2004) and some of the world’s longest sea turtle monitoring programmes. Hawksbill turtles are tagged as part of monitoring (Allen *et al.,* 2010), allowing mothers to be identified. Where possible, we collected information on parent/clutch identity, lay dates, unhatched egg collection dates, and fates of other eggs in the clutch. At the end of incubation (∼60 days after oviposition), we opened and visually assessed the contents of failed eggs during routine nest excavations, and randomly collected one or two eggs per clutch that showed no signs of development. Egg yolks were stored in either 5-10% formalin, depending on the availability of different formalin concentrations at different field sites, and transported to the University of Sheffield where they were examined up to five months after collection. Of the 58 eggs received, 42 were examined from 35 different clutches. Insufficient material was retrieved from the other 16 eggs to allow examination.

#### Detecting PVM-bound sperm and embryonic nuclei in the germinal disc

Captive eggs were opened carefully by cutting around their shells with fine scissors; yolks from wild eggs were removed from formalin solution. Egg contents were checked for signs of embryonic development and, if possible, pieces of the PVM were removed from directly above the germinal disc using forceps and small dissecting scissors, following the methods of Birkhead *et al*. (2008) and Croyle *et al*. (2016). Germinal disc material, if visible, was siphoned from the yolk surface with a micropipette. If the germinal disc couldn’t be seen or the egg was degraded/infected, as much PVM as possible was extracted from the egg contents to maximise the chance of detecting embryonic cells microscopically. The PVM was washed in phosphate buffered saline (PBS) to remove excess albumin and yolk and placed, along with any germinal disc material, on a microscope slide. A nucleic acid dye, Hoechst 33342 (0.05 mg/ml), was then applied, followed by a coverslip, and slides were left for at least 10 minutes in the dark before examination (Croyle *et al*., 2016)

PVM and germinal disc material were examined at 100X-400X magnification using a fluorescence microscope with UV illumination, a BP 340-380 excitation filter, and LP 425 suppression filter (Birkhead *et al*., 2008). Photographs were taken for documentation (Figure 4). Care was taken to distinguish between sperm heads and microbes in microbially infected eggs, as Hoechst 33342 also stains fungal and bacterial DNA (Croyle *et al*., 2016; Phillott and Godfrey, 2020).

**Figure 4.**
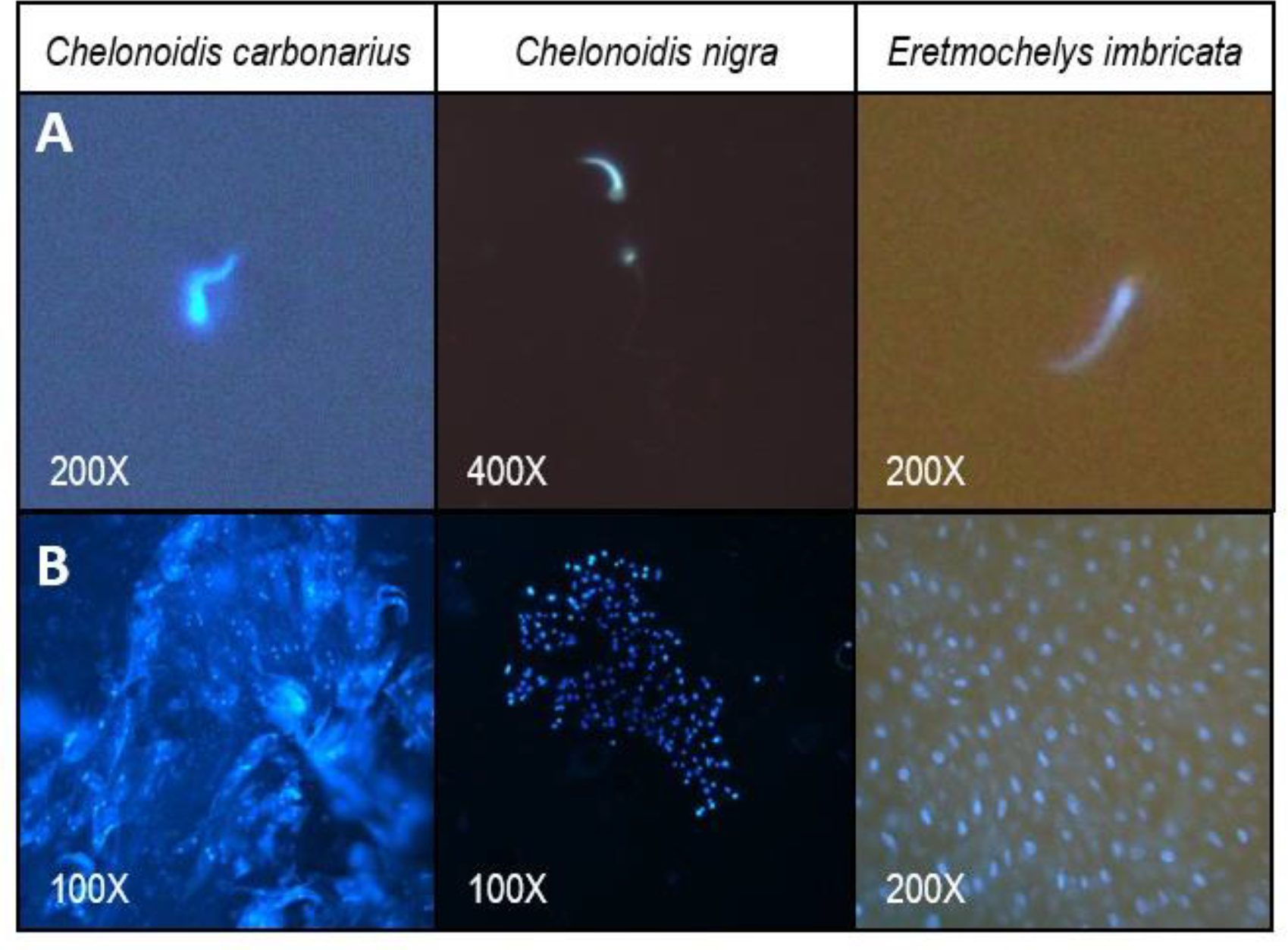
*Stained nuclei of **(Row A)** PVM-bound sperm, and **(Row B)** embryonic cells and/or tissue and the respective microscopic magnification levels for the Red-footed tortoise (*C. carbonarius – nuclei on perivitelline membrane*), Galapagos tortoise (*C. nigra*), and Hawksbill turtle (*E. imbricata).

Examined eggs were classified as fertilised if embryonic nuclei were identified in the germinal disc or adhered to the PVM. Eggs were classified as unfertilised if PVM from most/all of the yolk was retrieved and clearly observable under the microscope, and very few/no sperm and no embryonic nuclei were found consistently across all pieces of PVM. If there was insufficient PVM to be confident of fertility status (i.e., abundant embryonic nuclei could not be found), the egg was classified as inconclusive, even if PVM-bound sperm were detected. However, it should be noted that presence of PVM-bound sperm alone provides useful confirmation of male fertility/sperm availability.

#### Animal ethics and permits

This research was reviewed and approved by BIAZA to be carried out with UK zoo members, and by the Seychelles Bureau of Standards (SBS) to be carried out with Seychelles conservation organisations (Ref: A0157). Non-viable eggs were received opportunistically from these UK and Seychelles-based collaborators. Hawksbill turtle (*Eretmochelys imbricata*) eggs from Seychelles were authorised for export as per the agreement made with the Ministry of Agriculture, Climate Change and Environment, in accordance with Article 15 of the Convention of Biological Diversity. Additionally, permits were acquired for all eggs collected from species listed under the Convention of International Trade in Endangered Species of Wild Fauna and Flora (CITES; Seychelles export permit #A1517; UK import permit #615170/01)

#### Part 2: Results

We identified PVM-bound sperm and embryonic nuclei in three out of four species examined: the captive Red-footed tortoise and Galapagos tortoise, and the wild Hawksbill turtle (Figure 4). The single Spiny turtle egg that we obtained had a significant microbial infection which precluded its examination. However, microbial infection did not always prevent examination: we confidently identified sperm and/or embryonic nuclei in several infected eggs from the other species, aided by the fact that sperm heads and fungal/bacterial cells are morphologically distinct and easily discriminated (see Appendix B to compare with an example of microbial nuclei). Microbial infections were more common in refrigerated eggs compared to eggs that were frozen or formalin-preserved, and overall, wild Hawksbill turtle eggs fixed in 5% formalin were best preserved and proved easiest to examine. We were able to detect embryonic nuclei and PVM-bound sperm in undeveloped Hawksbill turtle eggs that had been left in nests for ∼60 days after oviposition followed by another <5 months preserved in formalin.

Overall, egg fertility status was determined conclusively in 70% of all microscopic examinations (Figure 5), and of these, only one egg – collected from a wild Hawksbill turtle on Cousin Island – was found to be unfertilised (Figure 5). All eggs with blood spots (Table 1) were identified as fertilised and, as expected, eggs with visible embryos and/or eggshell chalking (n = 14) consistently tested positive for the presence of embryonic nuclei, suggesting that these indicators are unlikely to generate false positive fertility assignments.

**Figure 5.**
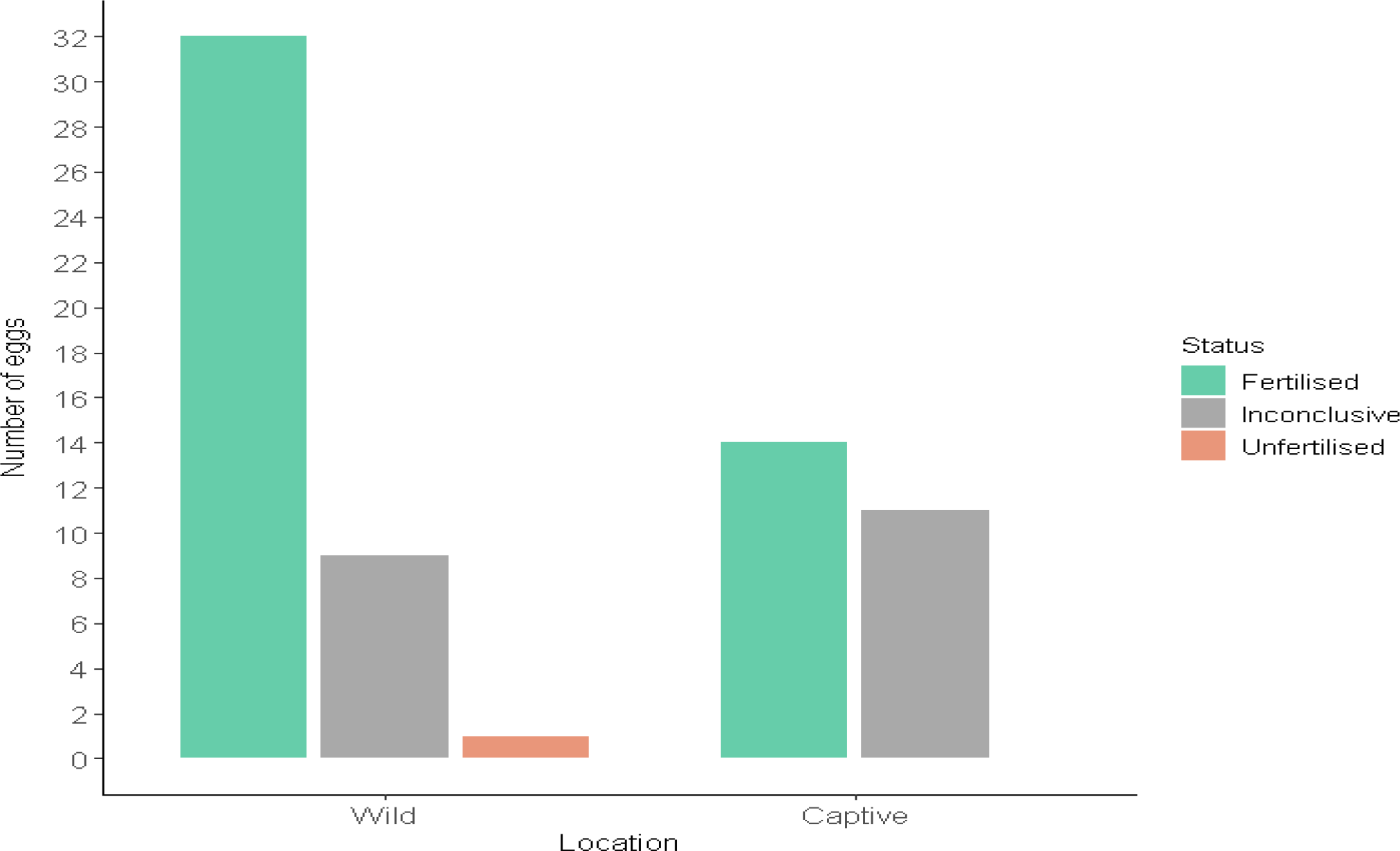
*The fertility status of 67 undeveloped eggs and those with blood spots (n = 4, all from UK zoo) were examined from captive populations from a UK zoo (Crocodiles of the World; n = 25) and wild populations from three different Seychelles islands (n =42): D’Arros (n = 7), Fregate (n = 9) and Cousin (n = 26). A total of 47 conclusive statuses were assigned (i.e., 70% of examined eggs) and 20 eggs remained inconclusive (30% of examined eggs). All eggs with blood spots were fertilised.*

## 6. Discussion

Here we have shown that current understanding of the relative roles of fertilisation failure and early embryo death in causing hatching failure in turtles and tortoises (Order Testudines) is limited, taxonomically biased, and suffers from methodological flaws. Most studies do not differentiate between fertilisation failure and embryo mortality, and when they do, the methods employed are inaccurate and/or unclear. Despite recommendations to use alternative methods (Phillott and Godfrey, 2020), studies have yet to adopt new techniques or test their efficacy across species/contexts. We have therefore attempted to move the field beyond this barrier by demonstrating the successful application of a new method across three species, including both captive and wild populations, and showing that they can be used to identify fertilisation success and sperm availability.

Most studies investigating turtle and tortoise hatching failure have not discriminated between fertilisation failure and embryo death as the cause, and as a consequence, our ability to fully understand the mechanisms underpinning early reproductive failure is limited. For example, Sinaei and Bolouki (2017) suggest that hatching failure in green sea turtles (*Chelonia mydas*) in Oman is associated with maternally transferred heavy metals. However, since the researchers did not investigate the fertility status of unhatched eggs, it remains unknown whether heavy metals primarily impact adult fertility (i.e., gamete production/quality), resulting in unfertilised eggs, or embryo development/survival via contaminated egg contents. The method we have described here may help address this knowledge gap in future investigations.

We have demonstrated that microscopic detection of PVM-bound sperm and embryonic nuclei in the germinal disc is usually (70%) successful and can provide conclusive evidence of fertilisation, even in eggs that have passed their incubation time and undergone a degree of degradation. The vast majority of successfully examined undeveloped eggs in our study were found to be fertilised (66/67 or 99%), indicating that traditional approaches significantly overestimate rates of fertilisation failure. Although all examined eggs with blood spots were fertilised, our sample size was small (n = 4) and further investigation is required to determine whether blood spots are a reliable indication of fertilisation.

Our literature review revealed biases in the study of hatching failure across turtles and tortoises (Testudines Order). Marine turtles are over-represented (especially *Caretta caretta*), tortoises are severely under-researched, and few studies consider captive populations or wild/captive comparisons. The bias towards wild populations is perhaps not unexpected; few captive breeding programmes exist for marine turtles due to the maintenance of healthy wild populations being prioritised, as well as species-specific challenges of maintaining sea turtles in captive conditions (Owens and Blanvillain, 2013). Wild marine turtle populations are also relatively easy to monitor, as their nesting seasons and locations are fairly predictable. In contrast, freshwater turtles and tortoises occur at lower densities, are more cryptic, and have limited seasonal and daily activities (Zylstra *et al*., 2010), making them more challenging to study. The focus on wild marine turtles may also explain why visual assessment of egg contents (Figure 2A) has been most commonly used to assess egg fertility – this is the most practical method in the field. Indeed, several studies explicitly state that they didn’t investigate egg fertility due to difficulties with accurately classifying undeveloped eggs (e.g., Pintus *et al*., 2009; Van Lohuizen *et al*., 2016), while others included a disclaimer about the accuracy of their methods (e.g., Gane *et al*., 2020a, 2020b). This highlights the need for a practical solution for both field and captive application.

Our findings indicate that methods for examining unhatched eggs are applicable across three taxonomically diverse species, and in both captive and wild contexts. Consistent with Croyle *et al.,* (2016), we found identifying the germinal disc to be difficult (more so than in birds; NH pers. obs.), yet we were still able to detect embryonic cells in 70% of examined eggs. Moreover, while finding only PVM-bound sperm is not conclusive evidence of fertilisation, it is nonetheless indicative of successful copulation and sperm availability – if few/no sperm are found, this may reflect issues with sperm production or transfer (Croyle et al., 2016).

The rarity of unfertilised eggs in our study suggests that reproductive behaviour and copulation problems, insufficient/defective sperm, or oviductal-sperm incompatibility (Birkhead *et al*., 2008; Hemmings *et al*., 2012), are unlikely to be a major problem for the wild Hawksbill turtle population nesting on the Seychelles Islands of Cousin, D’Arros and Frégate (note that findings are biased towards Cousin Island due to larger sample size). However, one Hawksbill turtle egg was found to be unfertilised: a triple-yolked egg from an unusual nest on Cousin Island. The clutch presented several issues; 83% of eggs showed no signs of embryonic development; eggs had irregular morphologies; and dwarfism was expressed in the few emerging hatchlings (Appendix C). All three yolks in our sample remained intact for laboratory investigation and the PVM of each was thoroughly inspected, consistently revealing no embryonic nuclei or sperm. Considering that (1) the examined egg was unfertilised; (2) most other eggs in the clutch were undeveloped; and (3) there were many irregular sized eggs, it seems likely that the relatively high rate of hatching failure in this clutch was linked to issues with the parents’ reproductive health.

We used a range of different egg storage methods prior to examination, including refrigeration, freezing, and formalin-fixation. This was largely due to logistical constraints, but nonetheless allowed us to make preliminary assessments of the suitability of different methods. Refrigeration was the least effective: compared to frozen of fixed eggs, refrigerated eggs were more likely to develop or progress existing microbial infections. This may explain why captive eggs (typically refrigerated) were generally more difficult to analyse than wild eggs (all formalin-fixed) (Figure 5). However, eggs from different species may react differently to different forms of preservation. For instance, in some Galapagos tortoise eggs, the yolk surface turned grey in colour after freezing, which made identifying the germinal disc impossible but did not affect microscopic examination of the PVM. This did not happen in frozen Red-footed tortoise eggs. Eggs appeared to be best preserved in 5% formalin; 10% formalin was also effective, but egg contents became somewhat more brittle and more challenging to dissect. Importantly, embryonic nuclei and PVM-bound sperm detection was possible in undeveloped eggs that remained in the nest for ∼60 days after oviposition, followed by <5 months stored in 5% or 10% formalin, addressing the concern raised by Phillott and Godfrey (2020) that the long incubation time (approximately 50 days) and nest cavity conditions of many turtle species may degrade failed eggs to the point that they cannot be examined.

In summary, we have identified important gaps in our understanding of turtle and tortoise (Order Testudines) hatching failure and provided new tools for monitoring egg fertility and embryo survival rates in threatened species. We recommend that future research combines accurate data on fertilisation failure and embryo mortality rates, generated using the approach we have described, with data on breeding conditions (e.g., nest site temperature), conservation interventions (e.g., nest relocations), and other potential drivers of reproductive failure such as pollutants and disease exposure, to monitor the impact of environmental change on early reproductive processes. We also anticipate that this approach will be applicable to other reptile groups, such as crocodiles (Augustine 2017), allowing scope to investigate similarities across birds and reptiles. The methods outlined here may therefore equip researchers and conservationists working across broad taxonomic groups with a tool to inform conservation and breeding management in the face of global change.

## Acknowledgements

We sincerely thank all people and organisations that made this study possible. The conservation teams and volunteers involved in sample collections from BIAZA members (specifically, Colin Stevenson (Head of Education) and the team at Crocodiles of the World, UK, and Dr Christopher Michaels (Herpetology Projects Manager) and the Herpetology team at ZSL Whipsnade Zoo), Nature Seychelles, Fregate Island Conservation Team, and D’Arros Research Centre team (specifically, Henriette Grimmel, Ellie Moulinie and Dillys Pouponeau) have been paramount. The incredible moral and logistic support by Mrs Sabrina Lapolla Victor and Miss Shireen Oliaji greatly assisted with the fluency on the project. Thanks to the BIAZA Research Committee for supporting the project and encouraging members to contribute samples. Lastly, Mr Elvis Nicette of SBS for approving this research to be carried out in the Seychelles. NH was supported by a Royal Society Dorothy Hodgkin Research Fellowship (DH160200). AL was supported by a fellowship enhancement award from the Royal Society to NH.

---

**Appendix A.** *Turtle and tortoise hatching success studies investigating the fertilisation status of unhatched eggs that have met the criteria of the systematic review. Their investigated Testudines species, the context of the study population (whether eggs were collected for wild or captive conditions) and the method(s) used to assess fertility of unhatched eggs, and necessary justifications are presented.*

| Citation | Species | Study population context | Method(s) of classifying egg fertility (i.e., both fertile and unfertilised eggs) |
| --- | --- | --- | --- |
| Abella *et al.* (2007) | Loggerhead Turtle (*Caretta caretta*) | Wild | Visual assessment of a white spot on the eggshell surface and candling |
| Abella *et al.* (2017) | Loggerhead Turtle (*Caretta caretta*) | Wild | Visual assessment of a white spot on the eggshell surface and candling |
| Al-Mansi *et al.* (1999) | Green Sea Turtle (*Chelonia mydas*) | Wild | Unclear: excavations took place, but method lacking description |
| Ayamak *et al.* (2020) | Green Sea Turtle (*Chelonia mydas*) | Wild | Unclear: “Undeveloped eggs...were identified according to Kaska and Downie (1999)” – who describes stages of development and provides no details regarding differentiation between fertilised and unfertilised unhatched eggs. |
| Behera *et al.* (2018) | Olive Ridley Turtle (*Lepidochelys olivacea*) | Wild | Unclear: excavations took place, but method lacking description |
| Bell *et al.*, (2003) | Leatherback Turtle (*Dermochelys coriacea*) | Wild | Visual assessment of a white spot on the eggshell surface and of egg contents |
| Bladow and Milton (2019) | Loggerhead Turtle (*Caretta caretta*)<br>Green Sea Turtle (*Chelonia mydas*) | Wild | Visual assessment of a white spot on the eggshell surface |
| Blanck and Sawyer, (1981) | Loggerhead Turtle (*Caretta caretta*) | Wild | Visual assessment of egg contents |
| Bourou *et al.*, (2001) | Ploughshare Tortoise (*Geochelone yniphora*) | Wild+Captive | Visual assessment of egg contents |
| Buhlmann and Osborn (2011) | Wood Turtle (*Glyptemys insculpta*) | Wild | Visual assessment of egg contents |
| De Andrés *et al.* (2016) | Leatherback Turtle (*Dermochelys coriacea*) | Wild | Visual assessment of egg contents |
| De Solla *et al.* (2003) | Eastern Spiny Softshell (*Apalone spiniferus spiniferus*) | Wild | Visual assessment of egg contents |
| Escalona and Fa (1998) | Terecay Turtle (*Podocnemis unifilis*) | Wild | Visual assessment of a white spot on the eggshell surface and of egg contents |
| Gane *et al.* (2020a) | Hawksbill Turtle (*Eretmochelys imbricata*) | Wild | Visual assessment of egg contents |
| Gane *et al.* (2020b) | Hawksbill Turtle (*Eretmochelys imbricata*) | Wild | Visual assessment of egg contents |
| Garrett *et al.* (2010) | Leatherback Turtle (*Dermochelys coriacea*) | Wild | Visual assessment of a white spot on the eggshell surface and of egg contents (according to Whitmore and Dutton, 1985) |
| Geist *et al.* (2015) | Western Pond Turtle (*Emys marmorata*) | Wild | Visual assessment of a white spot on the eggshell surface and candling |
| Gidiş and Kaska, (2004) | Nile Soft-shell Turtle (*Trionyx triunguis*) | Wild | Visual assessment of egg contents |
| Hitchins *et al.* (2004) | Hawksbill Turtle (*Eretmochelys imbricata*) | Wild | Visual assessment of egg contents |
| Hopkins *et al.* (2013) | Snapping Turtle (*Chelydra serpentina*) | Wild | Candling |
| Ilgaz and Baran (2001) | Green Sea Turtle (*Chelonia mydas*)<br>Loggerhead Turtle (*Caretta caretta*) | Wild | Unclear: excavations took place, but method lacking description |
| Ji *et al.* (2003) | Chinese Soft-shelled Turtle (*Pelodiscus sinensis*) | Captive | Visual assessment of a white spot on the eggshell surface |
| Ji-chao *et al.* (2011) | Keeled Box Turtle (*Cuora mouhotii*) | Wild | Visual assessment of egg contents |
| Karavas *et al.* (2005) | Loggerhead Turtle (*Caretta caretta*) | Wild | Unclear: excavations took place, but method lacking description. |
| Langer *et al.* (2020) | Spiny Softshell Turtle (*Apalone spinifera*)<br>Snapping Turtle (*Chelydra serpentina*)<br>Northern Map Turtle (*Graptemys geographica*)<br>Blanding's Turtle (*Emydoidea blandingii*)<br>Midland Painted Turtle (*Chrysemys picta marginata*) | Wild | Visual assessment of egg contents |
| Leshem *et al.* (1991) | Nile Soft-shelled Turtle (*Trionyx triunguis*) | Wild | Unclear: no methods provided |
| Ligon and Lovern, (2009) | Alligator Snapping Turtle (*Macrochelys temminckii*) | Captive | Unclear: no methods provided |
| Mahdy *et al.* (2020) | Loggerhead Turtle (*Caretta caretta*) | Wild | Unclear: no methods provided |
| Markle *et al.* (2021) | Blanding's Turtle (*Emydoidea blandingii*)<br>Spotted Turtle (*Clemmys guttata*)<br>Midland Painted Turtle (*Chrysemys picta marginata*) | Wild | Unclear: excavations took place, but method lacking description |
| Mifsud *et al.* (2018) | Loggerhead Turtle (*Caretta caretta*) | Wild | Unclear – data under egg 'fertility' were actually descriptions of developmental stages and irrelevant to fertility status |
| Nicholson *et al.* (2020) | Eastern Box Turtles (*Terrapene carolina carolina*) | Wild | Unclear: no methods provided |
| Olgun *et al.* (2016) | Loggerhead Turtle (*Caretta caretta*) | Wild | Unclear: excavations took place, but method lacking description |
| Özdemir and Türkozan (2006) | Green Sea Turtle (*Chelonia mydas*) | Wild | Visual assessment of egg contents |
| Pignati *et al.* (2013) | The Yellow-spotted River Turtle (*Podocnemis unifilis*) | Wild | Visual assessment of egg contents |
| Quesada-Rodríguez *et al.* (2021) | Leatherback Turtle (*Dermochelys coriacea*) | Wild | Unclear: no methods provided |
| Sari and Kaska (2017) | Loggerhead Turtle (*Caretta caretta*) | Wild | Visual assessment of egg contents |
| Simon (1975) | Green Sea Turtle (*Chelonia mydas*) | Wild | Unclear: no methods provided |
| Simon *et al.* (1975) | Green Sea Turtle (*Chelonia mydas*) | Captive | Unclear: no methods provided |
| Sparling *et al.* (2006) | Red-eared Sliders (*Trachemys scripta elegans*) | Captive | Visual assessment of a white spot on the eggshell surface |
| Stancyk *et al.* (1980) | Loggerhead Turtle (*Caretta caretta*) | Wild | Unclear: no methods provided |
| Türkozan *et al.* (2003) | Loggerhead Turtle (*Caretta caretta*) | Wild | Visual assessment of egg contents |
| Ulrich and Parkes (1978) | Green Sea Turtle (*Chelonia mydas*) | Captive | Visual assessment of egg contents |
| Wei *et al.* (2021) | Reeves' Turtle (*Mauremys reevesii*) | Captive | Visual assessment of a white spot on the eggshell surface |
| Whitmore and Dutton (1985) | Leatherback Turtle (*Dermochelys coriacea*) | Wild | Visual assessment of a white spot on the eggshell surface and of egg contents |
| Wyneken *et al.* (1988) | Loggerhead Turtle (*Caretta caretta*) | Wild | Visual assessment of egg contents |

**Appendix B.**
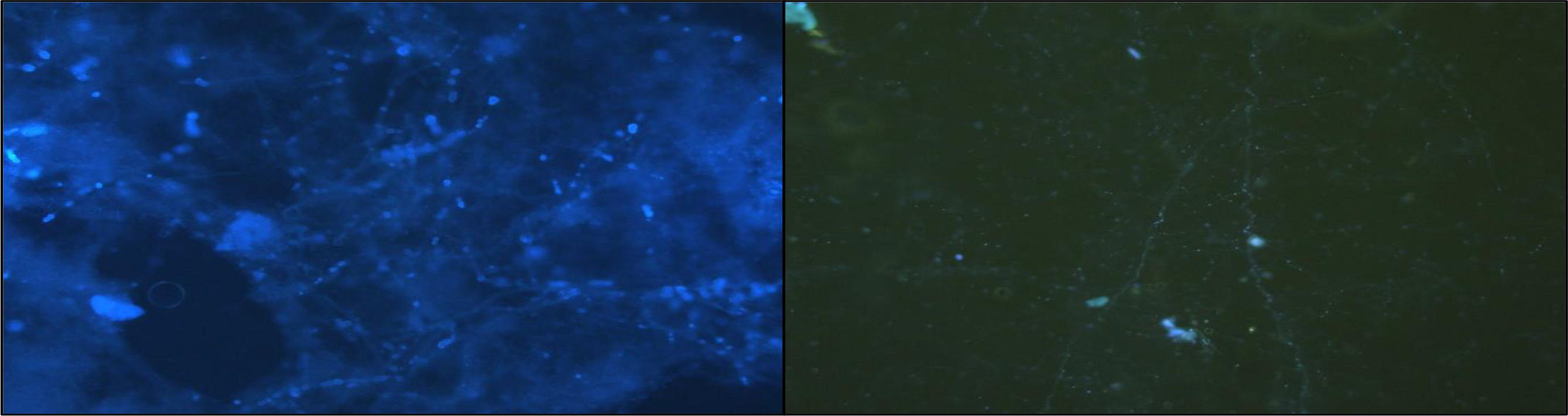
*Two 200X images of fungal/bacterial DNA found in the perivitelline membrane of a Spiny turtle (Heosemys spinosa) egg stained with Hoechest 3342.*

**Appendix C.**
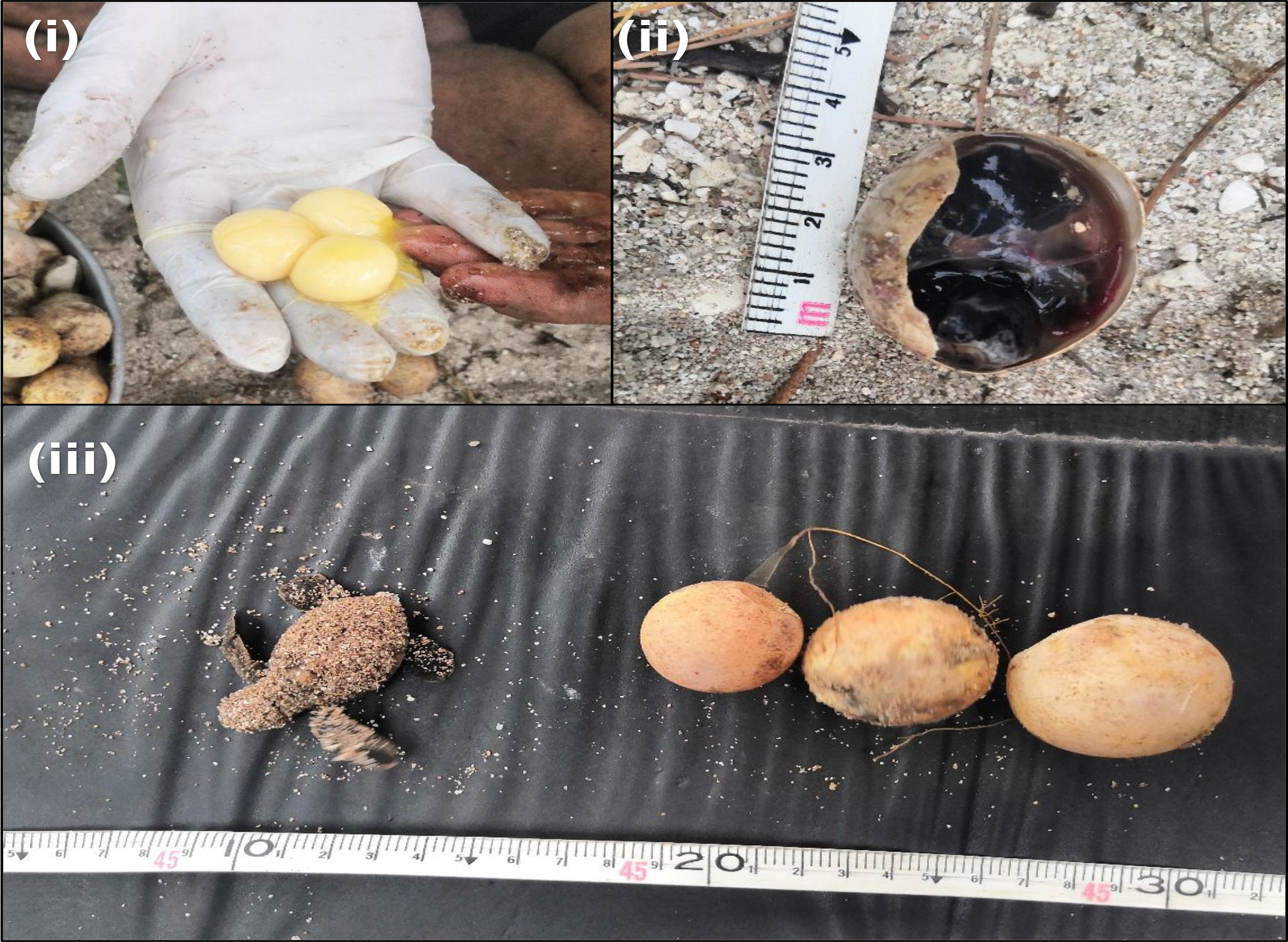
*An unusual Hawksbill turtle (Eretmochelys imbricata) egg clutch from Cousin Island, Seychelles that had several anomalies. **(i)** A triple yolked egg with no signs of embryonic development that was collected as a sample for lab investigation. Each yolk was smaller than an average sized yolk would be. **(ii)** Many eggs were unusually small, and some had unhatched developed embryos. **(iii)** A live dwarf hatchling alongside irregular sized eggs from the same clutch, ranging from abnormally small to large eggs*

